# Characterizing Subpopulations with Better Response to Treatment Using Observational Data – an Epilepsy Case Study

**DOI:** 10.1101/290585

**Authors:** Michal Ozery-Flato, Tal El-Hay, Ranit Aharonov, Naama Parush-Shear-Yashuv, Yaara Goldschmidt, Simon Borghs, Jane Chan, Nassim Haddad, Bosny Pierre-Louis, Linda Kalilani

## Abstract

Electronic health records and health insurance claims, providing observational data on millions of patients, offer great opportunities, and challenges, for population health studies. The objective of this study is identifying subpopulations that are likely to benefit from a given treatment using observational data. We refer to these subpopulations as “better responders” and focus on characterizing these using linear scores with a limited number of variables. Building upon well-established causal inference techniques for analyzing observational data, we propose two algorithms that generate such scores for identifying better responders, as well as methods for evaluating and comparing these scores. We applied our methodology to a large dataset of ~135,000 epilepsy patients derived from claims data. Out of this sample, 85,000 were used to characterize subpopulations with better response to next-generation (“Newer”) anti-epileptic drugs (AEDs), compared to an alternative treatment by first-generation (“Older”) AEDs. The remaining 50,000 epilepsy patients were then used to evaluate our scores. Our results demonstrate the ability of our scores to identify large subpopulations of epilepsy patients with significantly better response to newer AEDs.

## I. INTRODUCTION

Acentral problem in population health studies is inferring the influence of a treatment, or intervention, on a given outcome. For example, the influence of the drug “metformin” on the risk for cancer incidence [1]. There are several widely-used statistical techniques for estimating the average effect of a treatment, with respect to a given population [2]. However, the average effect may vary across different subpopulations, due to differences in individual susceptibilities to treatment. Therefore, certain patient groups may show stronger, or alternatively, weaker, response to treatment, than the larger population. In this study we focus on identifying “better responders”, which are subpopulations that are likely to benefit more from a given treatment, compared to the larger population.

Ideally, estimating whether a patient has a better response to a treatment requires comparing the outcome in two “parallel realities”; one in which the treatment is given, while in the other the alternative is used. We refer to these compared outcomes as “counterfactual”, as only one of them can be observed. Consequently, identifying better responders is essentially different from supervised learning problems, since we have no “better response” labels for the patients. In recent years there has been a growing interest in combining machine learning and principles of causal inference to construct models of *individual treatment effect* [3], [4]. However, these models are highly complex and do not provide interpretable characterizations of patients predicted as better responders.

Here we focus on finding interpretable models for identifying “better responders”, which point to the major factors that differentiates better responders from other patients. Such models may have a large applicability in Health Economics and Outcome Research (HEOR), as well as for building personalized treatment recommendation systems.

Validating a given characterization of better responders requires estimating the average causal effect of the treatment in the corresponding subpopulation. Ideally, average causal effects should be estimated with randomized controlled trials (RCTs), in which participants are randomly assigned to treatment groups. Unfortunately, RCTs are often costly, sometimes impractical, and may raise ethical questions. On the other hand, real world evidence (RWE) data, such as retrospective analysis of electronic health records and claims data, are abundant and offer new opportunities to study causal effects [5].

There are several common statistical methods for estimating average causal effects using observational data, such as inverse probability weighting (IPW) and standardization [2], [5]. These methods can be used to test specific hypotheses for better responder groups. One may suggest enumerating all hypotheses regarding the characterization of better responders and test each one using these validation methods. However, for high dimensional data, such as electronic health records, this approach implies a severe multiple testing problem and therefore becomes practically infeasible. Previous works presented methods for identifying better responders using randomized control trials [6], [7]. However, these studies are not directly applicable to observational data in which treatment assignment is not random. A different approach for identifying better responders suggest patients at high risk for the alternative treatment [8], [9]. However, such patients may not necessarily be at a lower risk with the treatment under consideration. More recently, decision trees were used to recursively partition the data into subpopulations that differ in the magnitude of their treatment effects [10] or by their optimal treatments [11]. In this study we take a different approach of learning sparse linear scores for better response. Such scores pinpoint the major factors associated with heterogenous treatment effects and enable the detection of subpopulations of various size showing better treatment effect using a score threshold.

In this work, we combine predictive modeling and causal inference theory to generate scores for identifying better responders using observational data. To ensure a simple and interpretable characterization of the identified subpopulations, we limit the generated scores to be sparse (i.e. including few variables) and linear. Building upon well-established techniques for estimating causal effects, we present two novel approaches for selecting the major variables that associate with differential response to the treatment. The generated scores are validated on a held-out test set, by verifying that the estimated average causal effect for groups of high-scored (respectively, low-scored) patients is larger (respectively, smaller) than the estimated average causal effect in the entire population.

We applied our methodology to a large dataset of epilepsy patients, comparing two alternative classes of AEDs: “Newer” vs. “Older”. The first class included second-generation AEDs that were approved over the last two decades for treating epilepsy in the US: felbamate, gabapentin, lacosamide, lamotrigine, levetiracetam, oxcarbazepine, pregabalin, tiagabine, topiramate, vigabatrin, and zonisamide. The second class contained the following first-generation AEDs that are available in the US market: carbamazepine, phenobarbital, phenytoin, primidone, and sodium valproate. In general, AEDs are initially approved as adjunctive therapy for patients with refractory epilepsy, based on data from randomized placebo-controlled trials. When an AED is initially marketed, there is uncertainty regarding the benefit to most epilepsy patients having less severe epilepsy and compared to the available Older AEDs. It is acknowledged within the clinical community that AED selection for epilepsy management should be optimized by adapting the treatment decisions to the characteristics of the individual [12]. Even when applied to a population that theoretically has a high chance to respond to it, prognosis remains difficult in most cases [13]. It is likely that there are characteristics that play an individual or interactive role in determining response / non-response that are currently unknown. This study aims to elucidate some of these characteristics by characterizing better responders to Newer AEDs using retrospective claims data.

## II. MATERIAL AND METHODS

We start by formulating the problem of characterizing better response, providing the necessary background and terminology on causal inference concepts (Section A). In Section B we present two algorithms for this problem. We present methods for evaluating and comparing scores for better responders in Section C. Finally, in Section D we describe the epilepsy use case in which we applied our methodology and describe specific implementation details. An overview of the entire methodology is presented in Fig. 1.

**Fig. 1.**
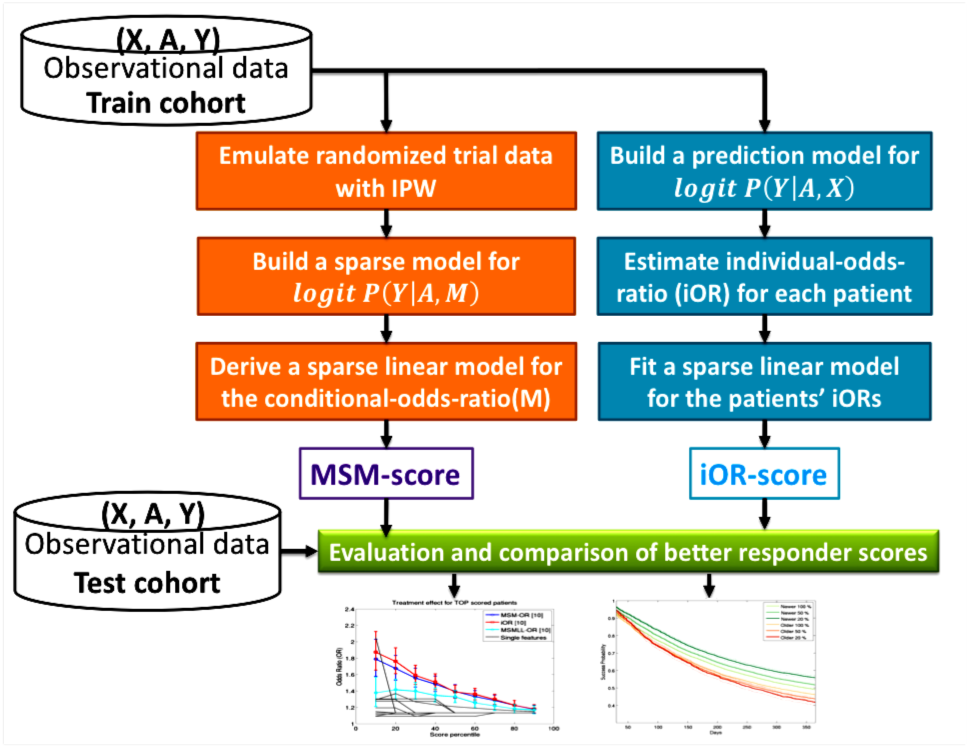
Overview of the methodology for generating and validating sparse linear scores for better responders. The symbols X, A, and Y correspond to observed variables, assigned treatment, and treatment outcome respectively.

### A. Problem definition

Suppose we have two treatment options, denoted by *a* = 1 and *a* = 0, and only one of these treatment options is given to a patient. Let *Y* be a random variable that indicates a patient outcome used for evaluating the response to the given treatment. Examples for outcomes may be: death, lab test result, hospitalization, etc. We denote by *A* the random variable that indicates the assigned treatment, and by *Y*^*a*^ the random variable corresponding to the potential outcome when *A = a.* The outcomes *Y*^*a*=0^ and *Y*^*a*=1^ are referred to as “counterfactuals”, as only one of them is observed for each individual: when *A = 1* then *Y = Y*^*a*=1^, and when *A = 0* then *Y = Y*^*a*=0^ is observed.

The average causal effect is defined by the *deviance* between the expected potential outcomes for the two treatment alternatives, *E*(*Y*^*a*=1^) and *E*(*Y*^*a*=0^). Specifically, when the outcome variable *Y* is dichotomous (e.g. hospitalized / non-hospitalized, death/survival) the average causal effect is the deviance between the two potential outcome *probabilities: p*_1_ = *P*(*Y*^*a*=1^ = 1), and *p*_0_ = *P*(*Y*^*a*=0^ = 1). Common measures for the average causal effect are the difference:

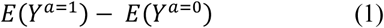
and the ratio:

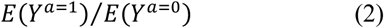
For a dichotomous outcome, it is also common to consider the *odds ratio* (OR) of *p*_1_ = *P*(*Y*^*a*=1^ = 1) and *p*_0_ = *p*(*y*^*a*=0^ = 1) as the measure of the effect:

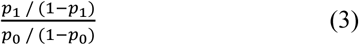
If the assignment of patients to treatments was random (i.e. A is randomly set), then *P*(*Y*^*a*^ = 1) could be estimated by *P*(*Y* = 1| *A* = *a*). Randomized trials use randomization of *A* for just this purpose. However, in observational data, such as electronic health records or claims data, treatment assignment (*A*) is usually far from being random and often depends on several factors that can potentially affect the outcome (*Y*). Such factors, which potentially affect both treatment assignment and the outcome, are referred to as *confounders.* The estimation of causal treatment effects from observational data must correct for biases in potential confounders.

Formally, to infer treatments effects from observational data we employ two assumptions. First, the standard *strong ignorability assumption* [14] that potential outcomes are independent of the treatment assignment when conditioned on the covariates *X* (i.e. no hidden confounders):

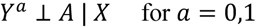
Second, we assume positivity:

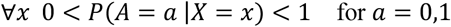
We refer to a subset of the observed variables *L* ⊂ *X* as a *sufficient set of confounders* if it satisfies the above conditions. Note that such set may not necessarily be unique. The expected potential outcomes, and consequently the average causal effect, may change between different subpopulations, e.g. men vs. women, older vs. younger. We say that a random variable *M* is an effect modifier when the average causal effect varies for subpopulations with different values of *M*.

We are now ready to formulate the objective of this study, which is to identify better responders using sparse linear scores. We assume the interpretation of the response to be monotonic and that a better response corresponds to higher values of the response. Let *X* be the set of observed variables, excluding the outcome variable *Y* and the treatment variable *A.* Given a relatively small number *k*, the goal is to find a subset {*M*_1_, …, *M*_*k*_,} ⊆ *X*, *k*’ ≤ *k*, and a linear score:

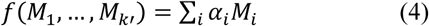
such that a subpopulation of patients with higher scores will have a better (i.e. larger) value for the average causal effect.

### B. The algorithms

We propose two different algorithms for generating sparse linear scores for better responders. Such scores highlight the major factors that differentiates better responder subgroups from the larger population, facilitating their characterizing. The two algorithms share several common properties: They are both receive as input: (i) *L*, a sufficient set of confounders and (ii) *X’* ⊆ *X* a set of potential effect modifiers; Additionally, each of the algorithms uses a stepwise variable selection on *X’* to generate a sparse linear model for the conditional average effect. The key difference between the two algorithms lies in the way they estimate the conditional average effects. The first algorithm learns a prediction model for the outcome, and uses it to estimate the expected causal effect for *each individual.* It then fits a sparse linear regression model to the estimated individual effects. The second algorithm estimates conditional average effects with linear marginal structural models (MSMs) [15]. These linear MSMs yield linear scores for conditional effects. The two algorithms are described in detail below.

#### The individual-effect score

For a sufficient set of confounders, *L*, we get *E*(*Y*^*a*^ | *L) = E(Y*|*A,L).* We assume that conditioning on additional variables from *X*, and in particular on the set of potential effect modifier *X’*, does introduce new biases. Therefore *E*(*Y*^*a*^|*L,X’) = E(Y*|*A,L,X’).* Suppose that we have a prediction model for *E*(*Y*| *A,L,X’)*, which generates predictions *Ê(Y*| *A,L,X’).* We can apply the model to each individual in the data and, and use the predicted values *Ê(Y*| *A = 1,L,X’)* and *Ê(Y*| *A = 0,L,X’)* as estimates for *E*(*Y*^*a*=1^ I *L,X’)* and *E*(*Y*^*a*=0^ | *L,X’).* This allows estimating the individual effect for each patient in the data, based on his/her own values for *L* and *X’.* Finally, we augment patients’ data with their estimated individual effects as labels, and fit a sparse linear model that approximates these labels. A formal presentation of this approach is given Algorithm 1.

#### The MSM-effect Score

MSMs are models for the average potential outcome *E*(*Y^a^)* [15], [5]. MSMs can be extended to include effect modifiers, that is, predict *E*(*Y^a^|M)* where M corresponds to one or more effect modifiers [15], [5]. MSMs use the IPW method to reweight the population, such that in the resulting pseudo-population the treatment assignment variable, *A*, is independent of the observed variables, *X.* Below we describe in detail the use of MSMs by the algorithm. Additional details on the implementation of the IPW method in the epilepsy case study are given in Section D.

Let *M* ⊆ *X’* be a set of effect modifiers. If the average causal effect is measured by the difference, *E*(*Y^a=1^) - E(Y^a=0^)*, then we use the following linear MSM:

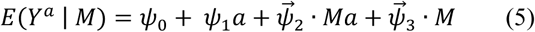
where 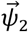 and 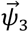 are coefficient vectors at the size of *M.* From this MSM we obtain a linear model for the conditional effect:

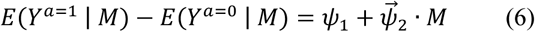
If the ratio *E*(*Y^a=1^)/E(Y^a=0^)* is used for measuring the effect, then we consider a linear MSM with log link function:

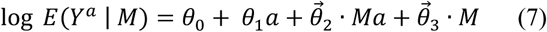
This MSM leads to a linear model for the *log* of the conditional effect

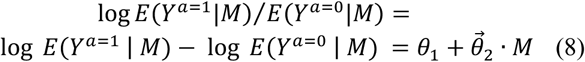
Finally, when using the odds-ratio for measuring the effect for a dichotomous outcome *Y*, we use the following linear MSM with logit link function

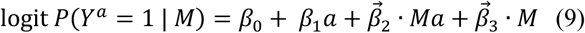
The odds-ratio measurement of the effect conditioned on M is:

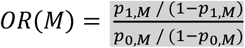
Where *p*_*a,M*_ = *P*(*Y*^*a*^ = 1 | *M*). We use the MSM in Eq. 9 to obtain a linear model for the *log* of the conditional effect

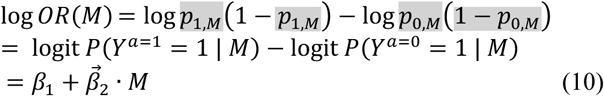
For each of the three causal effect measures that we consider: difference, ratio and odds-ratio, the corresponding linear MSM yields a linear score that estimates the conditional effect, or the log of it. Either way, the resulting score preserves the ranking of the patients induced by the conditional effect estimations predicted by the MSM. Consequently, higher scores correspond to subpopulations with larger estimated effect values.

#### Algorithm 1: Individual-effect Score

**Input:** *L* - a sufficient set of confounders, *X’* - potential effect modifiers, *k* - maximum number of variables

1. Fit a model 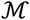 for predicting *E*(*Y*|*A,L,X’)*
2. For each individual *(L = l,X’ = x’):*
3. Use the model 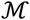 to predict potential outcomes *Ê(Y*^*a*=1^ | *L,X’) = Ê(Y*| *A = 1,L = l,X’ = x’)* and *Ê(Y*^*a*=0^ | *L = l,X’ = x’) = Ê(Y*| *A = 0,L = l,X’ = x’)*
4. Use predicted potential outcomes *Ê(Y*^*a*=1^ | *L = l,X = x’)* and *Ê(Y*^*a*=0^|*L = l,X’ = x’)* to estimate the individual effect
5. Fit a linear regression model 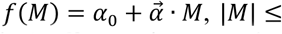 *k*, for predicting the individual effect, using stepwise selection on *X’*
6. **return** 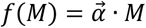

A variable is said to have *additive effect modification* if the corresponding coefficient in 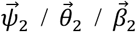 is significantly different than 0. The number of variables that we include in the MSM is limited, and hence we would like to select those having maximal additive effect modification. The MSM-effect score uses the greedy heuristic of stepwise variable selection, adding in each iteration a *pair* of terms *(x’ + x’a)* to the MSM, where *x’* ∈ *X’* has a maximal additive effect modification, if such exists. See Algorithm 2 for complete details on the MSM-effect score.

### C. Scores Evaluation and Comparison

The evaluation and comparison of the generated scores is done on a held-out test set. A score *f*(*M*) is expected to be an effect modifier since the average causal effect should vary across different levels this score. We verify that a score *f*(*M*) is an effect modifier by testing whether the corresponding random variable has an additive effect modification in the MSM *E*(*Y*^*a*^ | *f*(*M*)). To compare scores, we plot curves that map every score-percentile to the average causal effect computed within the corresponding group of individuals (i.e. top or bottom-scored individuals defined by that percentile). Ideally, higher score percentiles should correspond to larger average causal effect. For dichotomous outcomes that correspond to events, such as hospitalization and death, we analyze the corresponding time for these events. We use Kaplan-Meier curves to compare the distribution of *time-to-event* for the two potential outcomes, *Y*^*a*=0^ and *Y*^*a*=1^, corresponding to the treatment and its alternative. To account for the bias in treatment assignment, the Kaplan-Meier curves are computed on a reweighted population in which there is no bias between the two treatment groups (i.e. no confounders). We repeat this comparison for different scores percentiles, to verify that high-scored (respectively, low-scored) patients have larger (respectively, smaller) values for time-to-event.

#### Algorithm 2: MSM-effect Score

**Input:** *L* - a sufficient set of confounders, *X’* - potential effect modifiers, *k* - maximum number of variables

1. Use *L* to compute weights such that the reweighted population has no confounders
2. *M* ←∅
3. **Iterate** *k* times:
4. For each variable *X*_*t*_ ∈ *X*’:
5. Fit a linear MSM model with the set of variables *M* ∪ *{X_i_}* (see Equations 5, 7, 9 for difference, ratio and odd-ratio measures of causal effect)
6. Evaluate the additive effect modification of *X*_*i*_ in this MSM model using the P-value of the coefficient corresponding to the product term *X*_*i*_* *A* to be different from 0.
7. **If** at least one of the variables has an additive effect modification significantly different from 0:
8. *M ← M* ∪ *{X_imax_}* where *X*_*imax*_ is a variable having the most significant additive effect
9. **Else:** stop the iteration
10. **return** 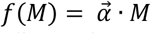, where 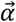 is the coefficient vector corresponding to the product terms in the final MSM. (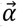 = 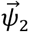 in Equation 5, 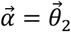 in Equation 7, 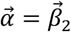 in Equation 9.)

We consider a variable selected to a score *f*(*M*) as robust, if it is likely to be selected on a different sample of the data. We test the robustness of the variables selected for each score using non-parametric bootstrapping. This is done by sampling the data with replacement, generating a new dataset of the same size as the original dataset (i.e. the same number of patients). We perform *Ν* iterations of bootstrapping, generating our scores for each dataset. The robustness of a selected variable is evaluated by the fraction of the times it is selected during *Ν* bootstrapping iterations.

### D. The epilepsy case study

#### The Data

To test our methodology, we used a dataset of ~135,000 epilepsy patients derived from the IMS Health Surveillance Data Incorporated (SDI) medical claims database. Epilepsy patients were identified based on their diagnoses and prescribed drugs. Every patient in our dataset was assigned with an index-date, which is the start of an AED treatment having exactly one added drug that was not in the previous AED regimen. We refer to this added drug as the *index drug.* The treatment starting at the index-date was classified as “Newer AED” *(A = 1)* or “Older AED” *(A = 0)* based on the class of the index drug. For more details on the data and study design see the appendix.

The year before the index-date is referred as the *baseline period.* It was used to derive variables *(X)* that characterize the patient in that time period. The year after the index-date is referred as the *treatment evaluation period* and was used to compute the outcome of the treatment *(Y).* See Fig. 2 for a schematic explanation of the referred time periods and their relation to variable computations. The evaluation of a treatment was done in the year following the index-date. Because the primary symptom of epilepsy - seizures - was not available in the SDI database, we used treatment changes as a proxy measure of seizure control and patients status. An unsuccessful outcome *(Y* = 0) was defined as any change other than a dose change (i.e. increase/decrease) or a complete withdrawal of any AED treatment in the subsequent 1 to 12 months after the index-date. A longer-term stable treatment or a complete withdrawal from an AED therapy was considered a successful outcome *(Y* = 1).

**Fig. 2.**
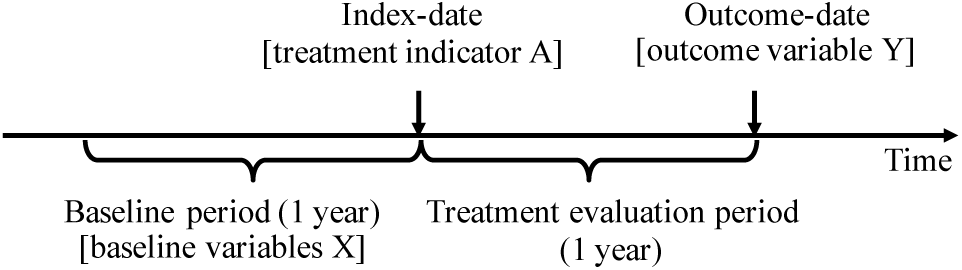
Overview of the methodology for generating and validating sparse linear scores for better responders. The symbols X, A, and Y correspond to observed variables, assigned treatment, and treatment outcome respectively.

The derived variables *(X)* included the following: age (at index date); gender; type of treatment change at index date; epileptic-state variables (indicator for each epilepsy-related ICD9 code, counts for each epilepsy-related Current Procedural Terminology (CPT) code, proxies for generalized/focal epilepsy as well as proxies for seizures [16]); medication possession ratio (MPR) and indicator for any use of AED and of each AED, indicator if patient used Newer/Older AEDs, the number of different AEDs, the number of treatment change events); comorbidities (based on an adapted list [17], as well as by diagnostic codes); mean monthly activity (using diagnoses, prescriptions and hospitalization data), indicator for hospital encounters, non-AED treatments (indicators for specific list of non-epilepsy drugs [17]), ecosystem variables (payer, state, first digit of zip code, specialty of physicians, year of index date).

Finally, the dataset was randomly partitioned into two sets: train and test datasets, which totaled ~85,000 and 50,000 patients, respectively.

#### The effect measure and scoring algorithms

Since we had a dichotomous outcome variable, we used the odds-ratio (OR) as a measure of the effect. We refer to the MSM-effect and individual-effect algorithms with the OR effect measure as MSM-OR and individual-OR respectively. We use the term iOR as an abbreviation for individual-OR. We generated the MSM-OR and iOR scores using the train dataset and evaluated and compared these scores on the test dataset.

#### Potential Confounders and Effect Modifiers

Identifying potential confounders is a key problem in causal analysis of observational studies [18]. In the epilepsy case study, we identified a set of potential confounders, *L*, using the following ad-hoc procedure. We first excluded nearly constant variables (mode frequency larger than 0.99). In accordance with a previous recommendation [19] and to avoid overfitting of our models for *Ρ (A | L)* and *P*(*Y | A,L)*, we included in *L* only variables that were significantly *(P < 0.05)* associated with *Y.* The association was measured by Chi-square (dichotomous variables) and t-test (continuous variables) after Bonferroni correction for multiple testing. Note that a variable is individually selected based on the strength of its association with *Y*, that is, without any reference to the P-values computed for the other variables. Finally, we filtered out from *L* variables that are highly correlated (Pearson correlation > 0.99) with each other, as such variables are expected to have low additive predictive value. We considered the resulting set of variables in *L* as a sufficient set of confounders. Since effect modifiers are also expected to be statistically associated with the outcome, *Y*, we used *L* as the set of potential effect modifiers and limited the stepwise selection procedure of the two algorithms, MSM-OR and iOR, to select variables only from *X’ = L.*

#### Outcome prediction model for iOR

For the outcome prediction model in the iOR algorithm (see step 1 in Algorithm 1) we used the following logistic regression model:

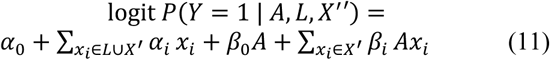
In general, other classifiers could be used as well for predicting the outcome, including random forest, SVM, gradient boosted trees, etc. Specifically, it is possible to make the prediction models more flexible by considering adding polynomial terms and additional interaction terms.

#### Generating balancing weights

In the epilepsy case study, we used the IPW method [2], [5] to generate balancing weights for the MSM-OR score (step 1 in Algorithm 2), as well as for evaluating and comparing the scores (Section II.C). The IPW method reweights every individual with assigned treatment *A = a* and observed confounder values *L = l*, with the following weight:

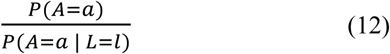
In this case study *P(A = a | L = l)* was estimated using a logistic regression model that was fitted to the data. Note that models for *P(A = a | L = l)* should be evaluated based on their ability to minimize the bias between treatment groups, and not based on their accuracy [2]. In the next section we describe a standard statistical method for evaluating the bias between treatment groups.

#### Testing imbalances between treatment groups

After generating a pseudo-population using IPW, we validated that all observed variables show no major imbalances between the two treatment groups. We quantified the balance for each variable using its standardized difference, *d*, which is the (absolute) difference in the variable means between the two treatment groups, divided by the combined standard deviation. To be exact, we used the following definition of the standardized difference,

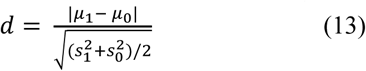
where *μ*_1_ and *μ*_0_ denote the average and *s*_1_ and *s*_0_ denote the sample variance of the variable in two treatment groups. Following [2], we considered a variable as balanced if its standardized difference was below 0.1.

## III. RESULTS

In this section, we present the results of applying our methodology to the dataset of epilepsy patients. We start by providing some descriptive statistics on the dataset.

### A. Data Statistics

Table 1 presents characteristics of the train and test datasets, demonstrating that the two datasets share the same data distributions. The causal effect of using a Newer AED in the entire population was estimated using the odds-ratio after balancing biases between treatment groups with IPW. The corrected OR values, which were independently computed in the train and test pseudo-populations, indicated that Newer AEDs had a positive casual effect on the outcome. As a comparison, the uncorrected OR values, which were computed in the original train and test datasets, erroneously indicated no causal effect. This striking difference between the correct and uncorrected OR values exemplifies the importance of correcting for the biases in treatment assignment *A.*

**TABLE 1.** TRAIN AND TEST DATA CHARACTERISTICS

| Characteristic | Train dataset | Test dataset |
| --- | --- | --- |
| Size | 83184 | 50000 |
| Index-drug is Newer (A=1) | 74 % | 74 % |
| Successful treatments (Y=1) | 49 % | 49 % |
| Gender, female | 62 % | 62 % |
| Age at index date (years) | 51±16 | 51±16 |
| Index-drug by neurologist | 42 % | 42 % |
| Index-treatment is monotherapy | 57 % | 57 % |
| Corrected odds ratio (OR) | 1.10 | 1.14 |
| 95% confidence interval | [1.07-1.14] | [1.09-1.18] |
| P-value | 1E-9 | 4E-10 |
| Uncorrected odds ratio (OR) | 0.98 | 0.99 |
| 95% confidence interval | [0.95-1.01] | [0.95-1.03] |
| P-value | >0.1 | >0.1 |

The total number of variables in *X* was 682. We selected the set of potential confounders by applying the methodology in Section II.D. We first excluded 308 nearly constant variables. Of the 374 remaining variables, 173 variables were found to be significantly associated with the outcome *Y* after Bonferroni correction. Finally, we excluded five additional variables due to high correlation with other variables. Overall, the set of potential confounders *L* included 168 variables. As described above, *L* was also used as the set of potential effect modifiers, *X’.*

### B. The Scores

We generated the MSM-OR and iOR scores for *Κ = 10* on the train dataset. Table 2 presents the variables selected by each of the scores, as well as their coefficients and the number of times they were selected in 10 bootstrap runs. The sets of variables selected by the 2 scores largely overlap, sharing 9/10 of the variables. Unless stated differently, all the variables in Table 2 were computed for the entire baseline period (one year). For example, the variable “Had neurological dx” indicates whether the patient had at least one diagnosis of neurological comorbidity during the year before the index-date.

**TABLE 2.** THE VARIABLES SELECTED FOR THE MSM-OR AND IOR SCORES.

| Selected variables | MSM-OR | iOR |
| --- | --- | --- |
| Index-date AED switched a previous AED | 0.3 (8 / 10) | 0.4 (10 / 10) |
| Had neurological dx | 0.3 (4 / 10) | 0.3 (7 / 10) |
| Was treated by a neurologist | 0.2 (6 / 10) | 0.2 (4 / 10) |
| Pregabalin MPR** | -- (5 / 10) | 0.1 (1 / 10) |
| Older AED MPR** | 0.1 (0 / 10) | -- (0 / 10) |
| Age | -- (1 / 10) | -0.1 (5 / 10) |
| Had a seizure proxy* in the previous month | 0.6 (10 / 10) | 0.7 (10 / 10) |
| Received Older AED | -0.3 (5 / 10) | -0.2 (2 / 10) |
| Had Medicare | -0.1 (2 / 10) | -- (4 / 10) |
| Had lipid metabolisms dx | -0.2 (4 / 10) | -0.2 (3 / 10) |
| Had back problem dx | -0.2 (6 / 10) | -0.2 (2 / 10) |
| Had trauma-related dx | -0.5 (0 / 10) | -0.4 (1 / 10) |

The first number in each column indicates the coefficient, while the second number (in brackets) contains the number of times the variable was selected in 10 bootstrap trials.
\* A seizure is inferred by a claim for ER, hospitalization or ambulance with epileptic/seizure primary or secondary diagnosis.
\*\* MPR = medication possession ratio

The variable “Had a seizure proxy in the previous month” was the strongest variable in the two scores. It had the largest coefficient and was shown to be most robust since it was selected by the two scores in all bootstrap runs. Another variable that was selected by the two scores and was found to be very robust was: “Index-drug switched a previous AED”.

### C. Scores Evaluation

We tested the additive effect modification of the variables corresponding to the two scores, as described in Section II.C. Both scores showed very significant additive effect modification, with P-values of 7e-36 and 5e-31 for the iOR and MSM-OR, respectively.

We also tested an intuitive variant of the MSM-OR score that used a more standard objective for a variable selection (Step 8 in Algorithm 2): maximizing the overall likelihood of *P*(*Y | A,M).* This variant of the MSM-OR score, which we noted MSMLL-OR, also showed a significant additive effect modification with a P-value of 7e-17.

Fig. 3 compares the iOR and MSM-OR scores by plotting the average causal effect, measured by the OR, in top- and bottom-scored patient groups as a function of the score percentile used to identify these groups. As shown in Fig. 3 the iOR and MSM-OR scores had very similar performance. They both managed to identify large subpopulations of patients that have significantly larger, or smaller, OR values compared to the OR observed in the entire population. On the other hand, the MSMLL-OR was less successful in identifying groups with significant higher, or lower, OR. We tested the variables in Table 2 as single-variable scores and compared them to the previous multi-variable scores. As can be seen in Fig. 3, the gray lines that correspond to single-variable scores were not able to significantly identify better treatment-responders - with the expectation of the variable “Had a seizure proxy in the previous month”. Since this variable is dichotomous, it was able to identify one group of better treatment responders, which included 11% of the patients.

**Fig. 3.**
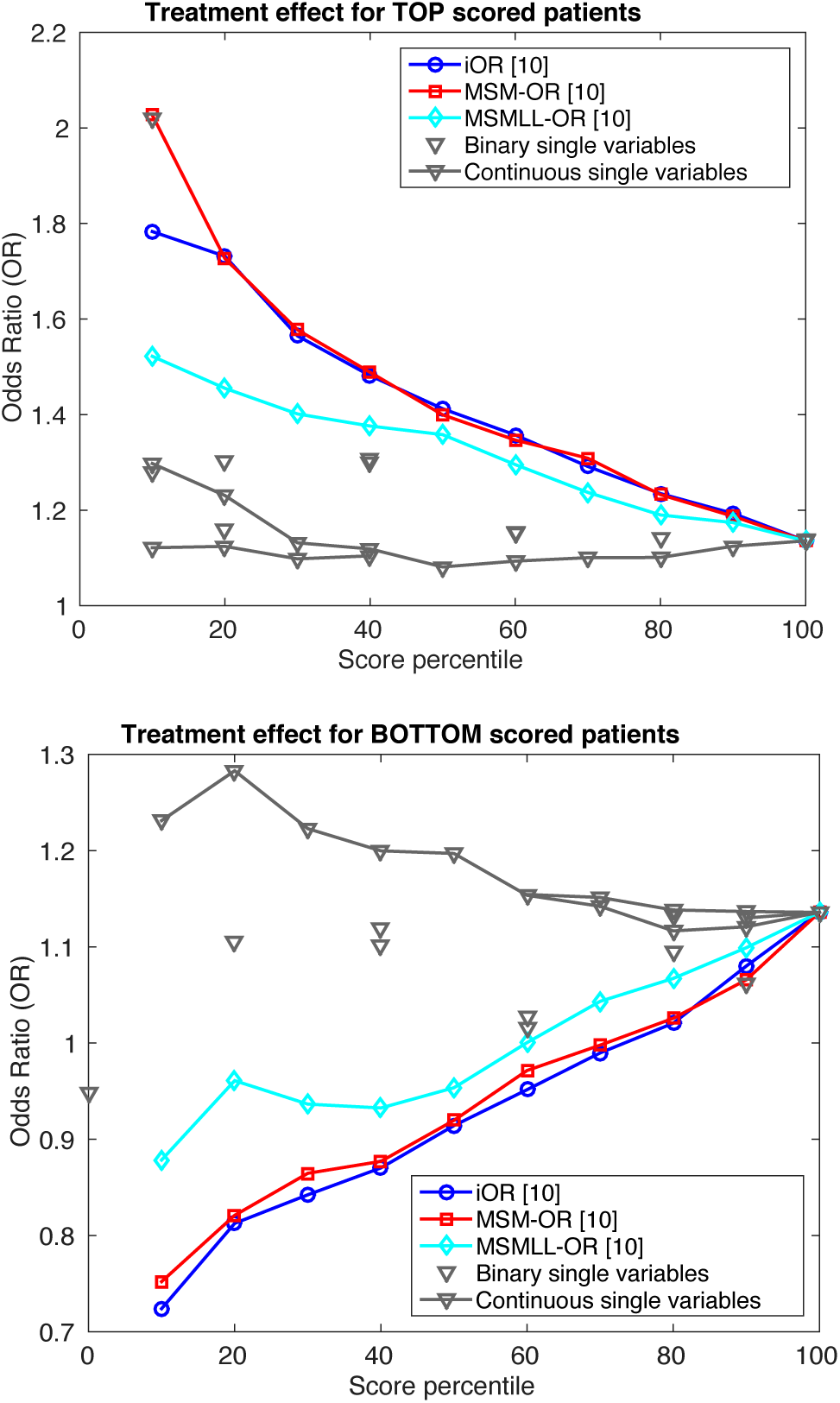
Scores evaluation and comparison. Gray lines refer to single-variable scores corresponding to the 11 variables that were selected to the MSM-OR and/or iOR scores.

We compared the time-to-failure in different score-groups, where a failure corresponded to a treatment change. Fig. 4 presents IPW-corrected Kaplan-Meier curves for the top-20%, top-50%, and top-100% (i.e. entire population) score-groups for the iOR score. As expected, the time-to-failure was longer on average for Newer AEDs in these score groups; this difference increased for score-groups with higher scores. We repeated the same analysis for bottom-score groups as well as for the MSM-OR score. In accordance to Fig. 3, the difference between Older and Newer in bottom-scored groups was in the expected direction (those on Newer AEDs having shorter time-to-failure than those on Older AEDs) but was less pronounced than the difference between top-scores patients (results not shown).

**Fig. 4.**
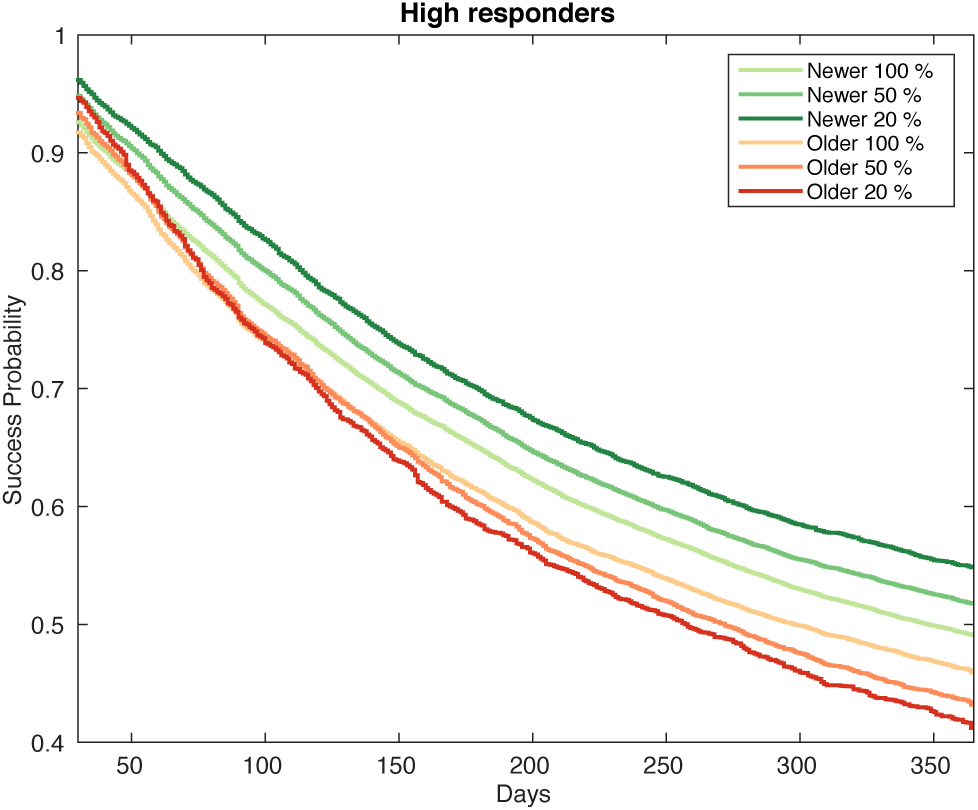
A comparison of Kaplan-Meier curves of time-to-failure for Newer and Older AED in top-scored patients by the iOR score. Newer/Older 20% means those on Newer/Older AEDs from the top-20% score groups and analogously for the 50% and 100% lines.

## IV. DISCUSSION

In the epilepsy case study, the two algorithms we presented, iOR and MSM-OR, yielded similar results both in terms of the set of the selected variables (Table 2), as well as in the ability to identify subgroups of better /worse responders in a held-out dataset (Fig. 3). In general, the two algorithms are *not* a-priori guaranteed to produce similar results, as they learn inherently different models of the data: the iOR algorithm model *P*(*Y | A, L)* while the MSM-OR algorithm model *P(A | L).* Thus, an agreement between the algorithms’ results strengthens their validity and the confidence in the underlying models.

Another major difference between the iOR/individual-effect and the MSM-OR/MSM-effect algorithms lies in the intricacy of the iterative variable selection procedure, which dominates the running times of the algorithms. The MSM-effect score uses a less-ordinary variable selection procedure, which in each iteration adds a *pair* of terms to a *generalized linear regression* (GLM) model. Conversely, the individual-effect score uses a standard variable selection procedure that in each iteration adds a *single* term to a *linear regression* model. Note that for the individual-effect score, the variable selection procedure could have been replaced by other variable selection methods, such as Lasso [20]. Fitting a linear regression model has a simple closed-form solution, which can be implemented in *0(nd^2^)* operations, where *η* is the number of samples and *d* is the number of variables [21]. In contrast, fitting generalized linear models with log or logit link functions involves an iterative method that maximizes the likelihood function [22]. Fitting generalized linear models in Matlab (glmfit function) and in R (glm function) is implemented with the iteratively reweighted least squares (IRLS) method, which takes *0(nd^2^)* operations *per iteration* [23]. The number of iterations for fitting each model depends on the convergence rate for the data. Overall, in our epilepsy case study, the MSM-OR score was three-time slower than the iOR score (results not shown).

In the epilepsy case study, the iOR and MSM-OR scores were trained to include 10 variables. The variables selected by the scoring algorithms are used as predictors for the differential response to Newer AEDs. Inspection of the selected variables (Table 2) shows that most relate to epilepsy and usage of AEDs. The occurrence of seizures and the existence of comorbidities are known to affect AEDs response. Variables that have been less clearly described in the past, include age and the related variable ‘Had Medicare’. Since AEDs are also prescribed for pain relief, it is possible that the selection of the variables ‘Had back problem dx’, and ‘Had trauma-related dx’ is a result of a contamination of our dataset with patients who consume AEDs for pain management. This is a drawback of using claims data, which do not make an explicit link between prescriptions and the diagnoses/medical conditions for which they were subscribed.

The iOR and MSM-OR scores showed superior performance to the MSMLL-OR score. This suggests that the variables selected by the iOR and MSM-OR scores better predict differential response to Newer AEDs than the variables selected by the MSMLL-OR score. Recall that the objective of the variable selection procedure in MSMLL-OR was to maximize the likelihood of the outcome prediction model. This implies that in this case study, the major predictors for differential response differ from the major predictors for the outcome itself.

In comparison to single-variable scores, the iOR and MSM-OR scores were much more successful in identifying better and worse responders (Fig. 3). The binary variable “Had a seizure proxy in the previous month” identified a single strong group of better responders totaling ~10% of the patients. Conversely, our iOR and MSM-OR scores could identify much larger groups of better responders in various sizes, with up to ~50% of the patients. Another interesting point is the ability of our scores to identify *worse* responders, that is, patients that are more likely to benefit from Older AEDs. While Older AEDs had a negative effect in the entire dataset, our scores identified a group with ~10% of the patients for which the odds for success were significantly lower for Newer AEDs than for Older AEDs’ (OR = 0.77 [0.68-0.87], P-value=5e-05). We note that the significance of the effect for this identified group of worse responders is much less pronounced than the effect observed in groups identified as better-treatment responders. For example, the treatment effect measured in the group corresponding to the 10% top-iOR scores was OR=1.87 [1.65-2.12], P-value= 1e-22.

There are several limitations to our study. In general, observational data studies are limited by the possible existence of unobserved confounders and selection bias. The claims data we analyzed were missing important data relevant to our study, such as seizure frequency, etiology, genetic data and/or neurological test results. Selection bias may exist due to the choice of patients showing regular medical activity. This selection of patients was done to control for potential data gaps due to the open nature of the database. Another limitation in our methodology relates to the selection of potential confounders by their statistical association with the outcome, in accordance with previous recommendations [19]. It may be preferable to base the detection of confounders on domain knowledge and causal diagrams that describe cause and effect relationships between variables [24]. However, identifying such a set in the presence of high-dimensional data where domain knowledge cannot capture the complex structure of the system is a challenging task of practical importance.

## V. CONCLUSION

Building on well-established concepts and methods from causal inference and machine learning, we presented two algorithms for characterizing subpopulations with differential response to treatments using observational data: individual-effect and MSM-effect. The two algorithms are designed to select the major factors associated with differential response for the given treatments. The differential response by itself is not observed for individual patients, and therefore traditional variable selection techniques, which require labeled data, cannot be directly applied. The two algorithms take different approaches for this hurdle. The individual-effect algorithm augments the train dataset with estimates of the differential response for each patient, thus reducing the problem into a supervised prediction task. On the other hand, the MSM-effect utilizes the causal inference method of linear MSMs to derive linear scores with most significant effect modifiers. We evaluated the generated scores by employing a machine learning train/test paradigm, that is, we tested the ability of scores to identify better/worse responders on a held-out dataset.

The use of sparse linear scores, which are commonly used in the medical domain for risk prediction, facilitates the understanding of the major “risk factors” for a differential response and their contribution to it. In the epilepsy case study we also used a simple logistic regression for modeling *E*(*Y | A, L)*, in the individual-effect algorithm and *P(A| L)* in the MSM-effect algorithm. In principle, other more sophisticated prediction models, such as Bayesian Additive Regression Trees [25] or boosted regression [26] may be considered for these intermediate learning tasks. In the recent years, various methods were proposed for generating balancing weights directly without modeling *P(A = a | L = l)* (e.g. [27]-[31]). Such methods can be incorporated into our framework, instead of the IPW method. Note that selecting the best method for a causal inference task is a challenge by itself, as the ground truth is unknown. A common approach to address this challenge is to test and compare the different models using simulated data under various mechanisms for generating the potential outcomes [32]. An interesting future work is to adapt such simulations for testing and comparing sparse models for differential response.

## APPENDIX

For this study, we used the IMS Health SDI medical claims databases containing anonymous, aggregated claims data of ~21 million patients from all major regions of the US. Data consist of diagnostic records (Dx), prescription records (Rx) and hospitalization records (Hosp). SDI is constructed as a provider-centric open database, in that it collects all data from providers (e.g. pharmacy) and identifies and links patients within that data. As such, SDI may contain unknown data gaps for individual patients if they visit providers that are not covered. To increase data reliability, we only considered data stretches in which there was 80% continuous monthly eligibility (in 1-year windows) in any of the SDI pharmacy, physician, or hospital databases, and quarterly pharmacy eligibility. This study was designed as a standard retrospective observational cohort study. The SDI data span 7 years, from January 1st, 2006, up to September 30, 2012. The index date is defined as the first valid treatment change event in which only one drug (from the Older and Newer AED lists above) is added. The conditions for an event to be a valid index date are defined as:

- The patient has at least 1 year of data before and after the index date.
- The patient has at least 3 months of Rx eligibility before the index date.
- During the 1-year period post index date, the patient is on some AED for at least 50% of the days. This is designed to exclude patients who are not actively consuming AEDs.
- The treatment was unchanged during the 30-day period after the index date (to eliminate rescue medication in favor of chronic treatment).
- There cannot be a prescription for the index drug in the year pre-index date.

To capture data from patients with epilepsy rather than from patients receiving AEDs for other indications, a patient had to fulfill the below criteria to be included:

- Diagnoses criterion: At least one International Classification of Diseases, Ninth Revision (ICD-9) epilepsy diagnosis code 345.* or two seizure diagnosis codes (780.39) at any time in the data.
- Prescription criterion: At least one claim for an AED at any time in the data. This claim must be from a pharmacy with 80% stability (existence of monthly pharmacy claims data) over the entire data period.
- Overall AED criterion: throughout the patient record, the patient had to have at least one AED claim which is not gabapentin/pregabalin or there had to be at least one gabapentin/pregabalin prescription from a physician whose specialty is one of: Neurology, Clinical Neurophysiology, Child Neurology, Neurological Surgery.
- In addition, we focused on adults, ≥16 years of age at the beginning of data.
- To avoid patients in whom AEDs were prescribed for indications other than epilepsy, the specialty of the physician prescribing the index drug was not allowed to be related to pain management or surgery.

Patients who met all of the above but for whom no valid index date could be found, were excluded.

## ACKNOWLEDGMENT

We thank Chen Yanover for his helpful comments on the manuscript.

